# Where is the money? Dynamics in feedback processing and attention during spatial probabilistic learning

**DOI:** 10.1101/2021.10.29.466415

**Authors:** Celina Pütz, Berry van den Berg, Monicque M. Lorist

## Abstract

Learned feature-based stimulus-reward-associations can modulate behavior and the underlying neural processing of information. In our study, we investigated the neurocognitive mechanisms underlying learning of *spatial* stimulus-reward-associations. Participants performed a probabilistic spatial reward-learning task that required participants, within 40 trials, to learn which out of four locations on a computer screen yielded the most gain-feedback when chosen. Our behavioral findings show that participants learned to choose which location was most rewarding. Those findings were paralleled by significant amplitude differences in event-related potentials (ERPs) elicited by the presentation of loss and gain feedback; the amplitude of the feedback-related negativity (FRN) was more negative in response to loss feedback compared to gain feedback, but showed no modulation by trial-number. On the other hand, the late positive component (LPC), became larger in response to losses as the learning-set progressed, but smaller in response to gains. Additionally, immediately following feedback presentation, brain activity in the visual cortex - read out through alpha frequency oscillations measured over occipital sites - was predictive of the amplitude of the N2pc ERP component, a marker of spatial attention orienting, observed on the next trial. Taken together, we elucidated neurocognitive dynamics underlying feedback processing in spatial reward learning, and the subsequent effects that spatial stimulus-reward association learning have on spatial attention.

## Introduction

Our ability to optimize behavior based on rewarding outcomes from the environment is an essential part of navigating through everyday life. Stimuli in our environment that we associate with rewarding outcomes shape our behavior, for instance by capturing our attention, a process that has been termed value-based attentional capture (Anderson, 2016). Research on value-based attentional capture revealed that reward-associated stimulus features can modulate attention voluntarily or involuntarily, whether they are task relevant or not (Bourgeois et al., 2017). This is manifested behaviorally through faster response times (RTs) in selecting target items that carry reward-associated features (Kiss et al., 2009), and slower RTs when distractor items do so (Anderson et al., 2016). However, behavioral findings on the effects of *spatial* reward-associations on attentional capture are conflicting, with some showing that reward-associated locations in our visual field can capture our attention similarly to reward-associated features (Mine et al., 2021; Sisk et al., 2020; Anderson & Kim, 2018, Chelazzi et al., 2014) whereas others do not find an effect (Won & Leber, 2018).

Studies investigating value-based attentional capture train their participants first in the reward-association, before testing it in an unrelated subsequent experimental task (Sali et al., 2014), suggesting that learning plays an important role in the capture of attention. Here, we sought to investigate the behavioral and neural mechanisms underlying spatial stimulus-reward learning, and their subsequent link to the neural processing of spatial attention.

In general, learning stimulus-reward associations is guided by feedback following a behavior. In the brain, subcortical reward areas are involved in the processing of feedback outcome (Delgado et al., 2005). In addition, frontal areas were found to contribute to the monitoring of these reward outcomes and to decision-making with regard to whether or not stimulus-reward associations and future behavior needs to be adjusted in particular (Rushworth et al., 2011). Lastly, the sensory cortices show changes based on the sensory features of the reward-associated stimulus (van den Berg et al., 2019; Henschke et al., 2020; Schiffer et al., 2014; Folstein et al., 2013). The result of such a process ensures that if the rewarded stimulus is encountered again in the future, it is processed faster and more efficiently (Hickey et al., 2010), thereby leading to the reward-associated attentional capture reported in the literature on feature-based reward associations (Anderson, 2016).

Electroencephalography (EEG) and event-related potential (ERP) recordings allow us to study the temporal sequence of processes involved in feedback processing during spatial stimulus-reward learning. For example, peaking around 200 ms after feedback presentation, the frontocentral negative-polarity feedback-related negativity (FRN) component was found to be more negative in response to loss feedback compared to gain feedback (Miltner et al., 1997; Hajcak et al., 2006; Holroyd & Coles, 2002; Nieuwenhuis et al., 2004; for a review see: San Martín, 2012). Traditionally, the FRN has been regarded as a neural correlate of the reward-prediction error (RPE), reflecting the difference between expected versus actual reward outcomes (Sambrook & Goslin, 2015, Cohen et al., 2007; Nieuwenhuis et al 2004). The RPE has been argued to be an essential part of learning, as the comparison between expected and actual outcomes is what makes individuals adjust their behaviors to match the actual reward outcome (Schulz, 2016).

The FRN is followed by the late positive component (LPC), a positive-going wave, with maxima occurring at parieto-central electrode sites, which starts around 400 ms after stimulus presentation (Trimber & Luhmann, 2017; Muller-Gras et al., 2019). The LPC has originally been studied in the context of affective processing (Paller et al., 1995). In the context of stimulus-reward learning, LPC amplitudes were observed to be larger in response to losses than to gains (Trimber & Luhmann, 2017). Furthermore, LPC amplitudes were found to be sensitive to reward expectancy, with largest amplitudes elicited by unexpected losses (Trimber & Luhmann, 2017; Muller-Gras et al., 2019, Donaldson et al., 2016). In addition, LPC amplitudes in response to losses are found to be linked to subsequent behavioral adjustments, suggesting that the LPC plays a role in behavioral optimization based on feedback (San Martín et al., 2013; von Borries et al., 2013). Thus, the LPC is reflective of the affective-value processing of feedback, which incorporates active context-updating based on the feedback that was received in order to to optimize behaviour (Polich, 2007; Glazer et al., 2018).

As an end-result of the feedback processing cascade, neural plasticity changes can be observed in the sensory cortices (Henschke et al., 2020; Schiffer et al., 2014; Folstein et al., 2013). Alpha power in the EEG signal (8 – 14 Hz) was, for example, significantly lower when measured over stimulus-specific sensory cortex in latency ranges later than the LPC (van den Berg et al., 2019), and has been inversely linked to the fMRI BOLD signal (Goldman et al., 2002, Scheeringa et al., 2011), indicating that activity in the alpha band might be related to cortical arousal. Importantly, these observations occurred in the absence of any sensory stimulation, after the presentation of feedback. The functional interpretation of sensory activity post-feedback is not clear. One possible explanation is that feedback-related top-down processes influence activity in the sensory cortex, in order to prepare for the upcoming choice. Similar neural mechanisms are observed in the context of preparatory spatial attention. Here, top-down influence of higher order brain areas is thought to induce a preparatory bias in the visual cortex before the onset of a stimulus array (Hopfinger et al., 2000). For instance, alpha power is significantly lower on contralateral occipital channels relative to the location that is about to be attended (Worden et al., 2000). Ultimately, this leads to enhanced processing of the stimulus occurring in that location (Zhao et al., 2019). Yet, whether feedback-related brain activity in the sensory cortex underlies a similar mechanism than preparatory activity during spatial attention has yet to be determined.

Lastly, spatial attentional orienting can be detected neurally via the N2pc ERP (Luck & Hillyard, 1994), a neural reflection of spatial orienting of attention that starts within 250 ms after the presentation of a bilateral stimulus array. The N2pc is a lateralized component that is larger on contralateral electrode sites of the attended location, and that shows larger amplitudes if the attended target is associated with reward (Kiss et al., 2009). In addition, N2pc amplitudes are predicted by preparatory alpha oscillations during attentional selection tasks, and are accompanied by alpha oscillations relative to the choice that was made (Zhao et al., 2019, Bacigalupo & Luck, 2019). Thus, the N2pc reflects a neural correlate of attentional processing in space.

In the presents study, we sought to investigate whether reward could be coupled to a spatial location, using a probabilistic reward-learning task. Additionally, we examined how the learning of spatial stimulus-reward associations could modulate spatial attention on a behavioral and neural level. Of specific interest was the cascade of feedback-related processes, ranging from feedback detection to stimulus-specific preparation of the sensory cortices, and whether those are related to subsequent processing of visuospatial attention. We first of all expected that participants would learn the spatial stimulus-reward association, as red out by an increased probability of choosing the highest-gain associated location-stimulus as the learning-set progresses. In addition, we expected that RTs would become faster in response to choosing the set-winning location-stimulus, indicating faster information processing as the reward-association is learned.

Neurally, we expected to observe amplitude differences between loss and gain feedback trials in the FRN component, with more negative FRN amplitudes in response to loss feedback. We further expected that, if the FRN represents a neural RPE, the difference between loss and gain feedback would become larger with trial progression as reward expectancy increases with learning. As an indication of context-updating, we expected to observe more positive LPC amplitudes for losses relative to gains, and that LPC amplitudes reduce in size once the location-reward association has been learned because less context-updating is needed. Following the LPC, but not before the next choice has been made, we hypothesized to observe significantly lower alpha power on contralateral electrode sites relative to the upcoming location choice, indicating stimulus-specific preparatory cortical activity. If indeed lateralized alpha activity reflects a preparatory process modulating the excitability in the sensory cortex of the to-be-chosen stimulus we further expected that feedback-locked alpha power contralateral to the upcoming choice would be significantly related to the orientation of attention towards the location of choice, indexed by the N2pc amplitude on the upcoming trial relative to the choice made.

## Methods

### Participants

Thirty-six adults participated in the experiment, of which two had to be excluded because of misunderstandings in the task instruction, and another three because of too many EEG artefacts (> 30 % of artefact-trials). Thus, the final sample comprised 31 adults (*M*_*age*_= 24.2, SD = 3.6), of which 23 were female and 29 were right-handed. All participants reported normal or corrected-to-normal vision. Four subjects participated for course credit, whereas 27 participated received monetary compensation (7 €/hour). In addition, participants received a bonus that depended on their task performance. Written consent was obtained from all participants before the start of the experimental session. Study protocols were approved by the Ethical Committee of Psychology of the University of Groningen.

### Apparatus

Participants performed the experiment in a dimly lit, sound-attenuated experimental room. The task was created and presented using the software package ‘Presentation’ (Version 20.3, Neurobehavioral Systems, Inc., Albany, CA, www.neurobs.com) on a 27-inch monitor with a refresh-rate of 60 Hz. Participants made their responses using a gamepad (Logitech Rumblepad).

### Procedure

After application of the EEG cap, participants were seated in a chair at approximately 60 cm distance from the monitor and instructed to keep their eyes on the fixation point at the center of the screen. In addition, participants were asked to limit movement during the experiment. They subsequently performed a probabilistic reward-based spatial learning task, in which they had to find a location-stimulus with the highest associated gain-probability using trial-by-trial gain/loss feedback as a guide, in order to maximize their monetary gains. On each trial they also had to bet either high or low on the to-be-chosen location. If participants placed a high bet and received gain-feedback on that trial, the monetary reward was 90 eurocents, whereas high bets and loss-feedback yielded a monetary loss of 90 eurocents. Low bets in combination with gain-feedback yielded a monetary gain of 30 eurocents, and low bets followed by loss-feedback led to a monetary loss of 30 eurocents. An interim summary after each learning-set of 40 trials provided subjects with the sum of money they had earned so far. Participants were paid out at the end of the experiment with a maximum of 15 €, the exact amount depending on their monetary gain during the experiment.

### Probabilistic reward-based spatial learning task

The probabilistic reward-based spatial learning task consisted of nine learning-sets, 40 trials each, spread across three blocks, totaling to 360 trials for the entire experiment. Within each learning-set, four locations (defined as four quadrants on the monitor screen) had a probability of monetary gain assigned to them. These four gain-probabilities were predefined as 0.3, 0.45, 0.55, and 0.7, and were randomly assigned to the four locations at the start of each learning-set.

Each trial started with a bet screen, which was shown until a response was made or 2000 ms elapsed. The bet screen consisted of a central fixation point with the letters “H” (meaning high bet) and “L” (meaning low bet) placed 50 pixels above or below the fixation. The letters randomly switched position on each trial, and participants had to press the upper left bumper button of the gamepad to choose the upper letter or the lower left bumper button to choose the lower letter. Following a response, the chosen bet option was highlighted for 300 ms. In case the participant made no response, a screen with the letters “no response!” was shown for 300 ms instead.

The bet screens were followed by a fixation point which remained onscreen between 700 and 900 ms. Next, the location stimulus screen appeared, in which four circle-stimuli containing a gap were presented. The circles were 90 by 90 pixels wide and placed in each quadrant of the screen, at a distance of 50 pixels from the fixation point. The four circles differed with regard to the location of the gap (up, down, left, right), which was linked to a button response using the right button-set on the gamepad. The buttons were arranged in the way that there was an upper, lower, left, and right button corresponding to the upper, lower, left, and right opening of the circle. Participants were instructed to choose a location by pressing the button that corresponded to the opening of the circle that was placed in the quadrant they wanted to choose on that trial. The position arrangement of the opening of the circles changed randomly on each trial. As a result, chosen stimulus location and button presses were unrelated.

The location stimulus screen was presented for 2000 ms, followed by a choice screen highlighting the location stimulus that was chosen (300 ms). After another central fixation screen (the duration again randomly ranging between 700 to 900 ms), feedback was presented for 500 ms. The words “gain” or “loss” occurred in black font on either an orange or blue background. The combination of feedback valence and background color was counterbalanced across participants, that is half of the participants received gain feedback on a blue background and loss feedback on an orange background and the other half vice versa. The trial ended with a third fixation screen, with a duration randomly chosen between 1700 to 2000 ms. Participants were allowed to take short, self-timed breaks in-between experimental blocks.

### EEG recording and pre-processing

EEG was recorded using a 64-channel, custom-layout, equidistant electrode cap and a DC amplifier at a sampling rate of 512 Hz and referenced to a central electrode channel (channel 5Z). Four subjects were measured using a sampling rate of 500 Hz, and data of these subjects were up-sampled to 512 Hz during pre-processing. Pre-processing was done using the MATLAB EEGlab (Delorme & Makeig, 2004) and FieldTrip (Oostenveld et al., 2011, http://fieldtriptoolbox.org) toolboxes. The recorded data was re-referenced to the average reference and filtered offline using a non-causal band-pass filter (Hamming windowed finite response filter, low cut-off value of 0.01 Hz, high cut-off value of 30 Hz). To correct for eyeblinks and horizontal eye-movements, we performed an independent component analysis using the algorithm implemented in EEGlab and reconstructed the EEG data without components that reflected eyeblinks or horizontal eye-movements (a maximum of three components per subject – *M* = 2.2 components). In addition, we excluded trials that exceeded an amplitude threshold of 120 mV, and excluded subjects that had more than 30 % of the trials removed. Lastly, we extracted epochs from -500 ms prior to the onset of the stimulus presentation screen to 2000 ms post-stimulus presentation screen, and applied baseline correction to the data starting at – 200 to 0 ms prior to the onset of the stimulus screen. We applied the same procedure to the feedback-locked epochs, this time using the onset of the feedback screen instead.

### ERP Analysis

In line with previous work (van den Berg et al., 2019), our regions of interest (ROIs) for the N2pc analyses were left and right occipital electrode sites (corresponding to PO7, O1, O2, PO8 in the 10-20 system). We determined time window of interest (TOIs) by creating stimulus-locked contralateral-minus-ipsilateral relative to the location of choice grand averages over all conditions and selecting a time-window from 50 ms before and after the peak, resulting in a TOI of 220 ms to 320 ms. We averaged amplitudes values in the TOIs over the left and right ROIs separately and then calculated the contralateral-versus-ipsilateral amplitudes by subtracting amplitudes of ipsilateral-to-choice channel sites from contralateral ones, and then collapsing over left and right choices (Luck & Hillyard, 1994).

We chose a fronto-central region of interest (ROI) for the FRN (corresponding to Fz, FCz, FC2, and FC1 in the 10-20 system, based on previous work; van den Berg et al., 2019). The latency interval was determined by creating a feedback-locked grand averages over trials and selecting a time-window corresponding to 50 ms before and after the grand average peak, resulting in an interval between 240 to 340 ms post-feedback onset. Our LPC ROIs were parieto-central channels, approximating Pz, CPz, P1, and P2 in the 10-20 system, and we selected amplitude values between 400 to 800 ms post-feedback onset. LPC ROIs and TOIs were derived from the literature (Glazer et al., 2018; Pornpattananangkul & Nusslock, 2015).

### Time Frequency Analysis

Time frequency analyses of alpha band power were done using the FieldTrip MATLAB toolbox (Oostenveld et al., 2011, http://fieldtriptoolbox.org). Frequency decomposition was performed on average-referenced EEG data time-locked to the feedback-presentation using a multitaper method based on discrete prolate Slepian sequences. We performed the analysis on linearly spaced frequencies from 1 to 40 Hz with steps of 0.5 Hz, using taper window widths that increased by one cycle every 3 Hz, starting at 3 cycles for 1 to 3.5 Hz and ending with 15 cycles for 37-40 Hz. Window smoothing was defined as frequency multiplied by 0.4, leading to less frequency smoothing in lower frequencies compared to higher frequencies. A log_10_ conversion was performed on the resulting power spectra to correct for non-normality of power within each time-by-frequency bin. Next, we extracted average alpha band power (8 to 14 Hz) between 600 and 1200 ms post-feedback presentation, from left and right occipital electrode channels (corresponding to PO7, O1, O2, PO8 in the 10-20 system) separately in order to analyze lateralized-to-choice alpha power. The ROIs for the analysis were chosen to be the same as for the N2pc analyses, and the TOIs were based on previous work (van den Berg et al., 2019).

### Statistical Analyses

No-response trials and trials in which RTs were faster than 200 ms (i.e. fast guesses) were excluded from the analysis. A no-response trial was defined as trials in which the bet or stimulus response was missing. Two subjects had incomplete datasets because of misunderstandings of the experimental task (< 11 % of trials and < 23 % of trials excluded) but were nevertheless included in the analyses. We applied a mixed-modelling approach using the lme4 statistical package (Bates et al., 2015), and for post-hoc analyses, the package emmeans (Lenth, 2021) was used, within the software package *R* (R Core Team, 2020). We determined the inclusion of random slopes varying by subject via model comparison, by selecting the model with the lowest Akaike information criterion (AIC, Akaike, 1974). This process allows for selecting the best-fitted model by taking over- and underfitting into account. In case the AIC difference yielded inconclusive results, we also took the Bayesian information criterion (BIC, Schwarz, 1978) of each model into account during model section. All models that we used in our analyses are summarized in **Table 1**. Degrees of freedom were approximated using Satterthwaite’s method using the package lmerTest (Kuznetsova et al., 2017). All models reported here contained the random factor ‘subject’ as a varying intercept.

**Table 1.**
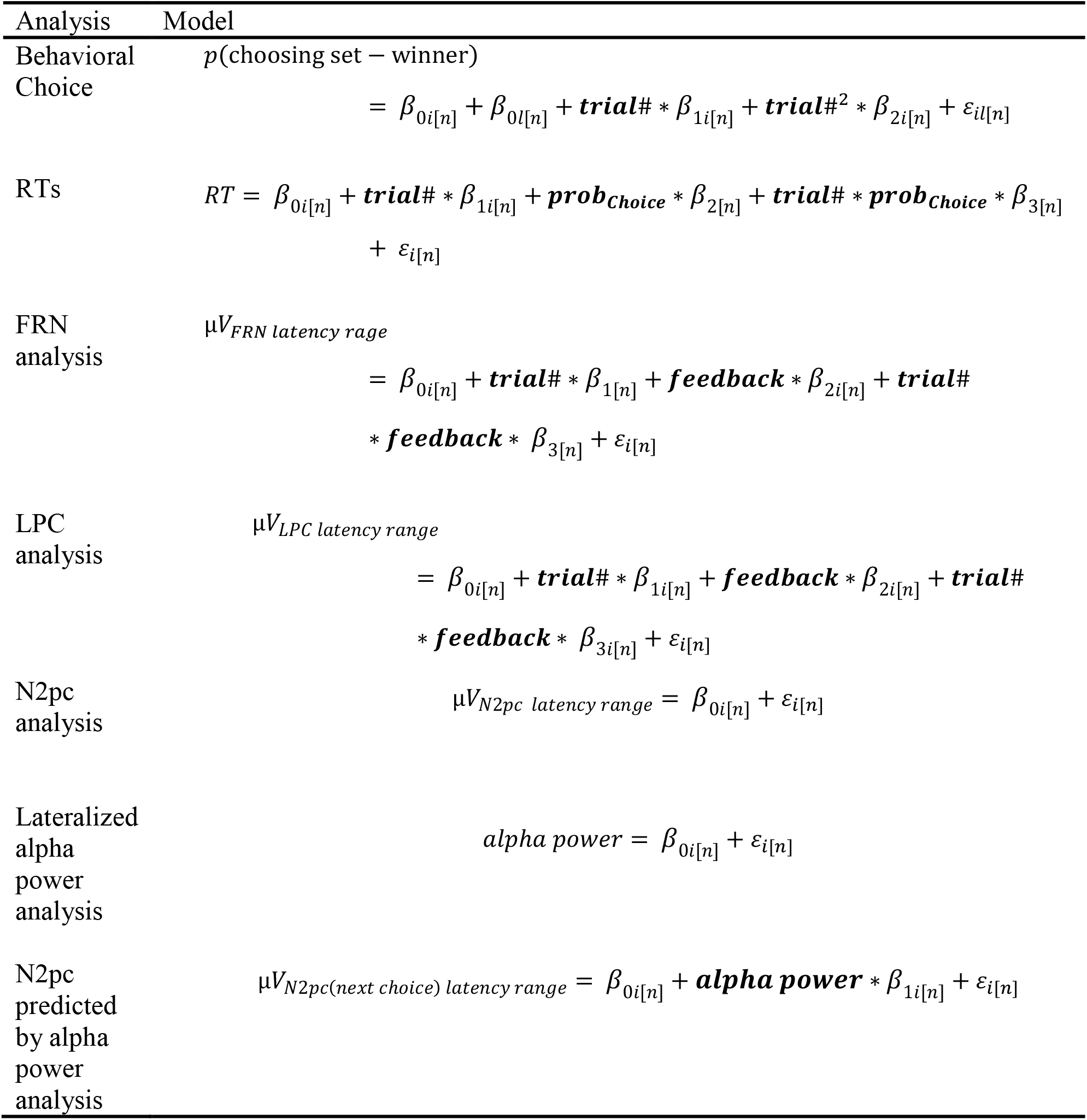
All models used in our analyses. Random effects (included based on AIC selection) are denoted by *i* and *l* by *n*.

### Behavior Analyses

We first tested whether participants showed a behavioral bias towards any of the locations by running a two-way ANOVA with the predictors ‘learning-set-half’ (with the two levels ‘first half of trials’ and ‘second half of trials’ of the learning-set) and ‘location’ (with four levels corresponding to the four locations). The predictor was the probability of choosing a location-stimulus when it was the assigned set-winner.

In order to analyze participant’s choice behavior with regard to the set-winner, we fitted a logistic mixed effects model with the binary outcome variable ‘choice: set-winner’ versus ‘choice: not set-winner’ as the dependent variable. As fixed effect predictors, we used the variable ‘trial-number’, ranging from 1 to 40, and the second-degree polynomial, to account for non-linearities in the data. The results of the behavioral model are tested with Wald’s Chi-Square test.

Initial observation of the RT data showed that responses made on the first trial of a set were substantially slower than all other trials (M_trial#1_= 904 ms, M_trial#1_= 696 ms, *t*(10804) = 15.6, *p* < 0.0001), and we therefore excluded the first trial from our model analysis. Next, we fitted a mixed-effects model with the outcome variable ‘Reaction Time’, and the fixed effect predictors ‘trial-number’, and ‘choosing set-winner’ versus ‘not choosing set-winner’, and an interaction between the two.

### EEG Analyses

Our analyses of the lateralized N2pc component relative to the location of choice included the predictor ‘trial-number’ and the random slope ‘subject over trial-number’. The FRN and LPC were analyzed by predicting their amplitudes with a fixed effect trial-number and feedback (gain and loss), as well as the interaction between trial-number and feedback.

### Time-Frequency Analysis

We analyzed average alpha power by first calculating the contralateral-versus-ipsilateral difference in power relative to the *next* choice, and excluded trials in which participants made a choice switch on the next trial. We fitted a linear mixed model with the N2pc amplitude as the dependent variable, and alpha power as the predictor.

## Results

### Behavior

#### Learning and Response Speed

The learning curves were not dependent on the set winning location, supporting that participants had no general bias towards one of the four stimulus-locations (interaction ‘learning-set-half by location’ *F*(3,188) = 0.57, *p* = 0.6; **Figure 1B**). In line with our expectations, the proportion of times that participants chose the set-winner compared to choosing a non-set-winner increased with increasing trial-number (Wald’s *X*^2^(2, 10806) = 204.07, *p* < 0.001, **Figure 2A**). Furthermore, the average amount of money that participants earned was 13.20 € (*SD* = 6.76 €), indicating that they indeed aimed at maximizing their monetary gains and learned the most rewarding location-stimulus. In addition, subject’s RTs decreased with increasing trial-number (main effect ‘trial-number’ *F*(2, 29.1)= 7.3, *p* < 0.01), and were significantly different between trials in which participants chose the set-winning location-stimulus versus when they chose a non-set-winning location-stimulus (main effect ‘choosing set-winner versus not choosing set-winner’ *F*(1, 10180.8)= 20.95, *p* < 0.0001; *M*_diff_= -17.51 ms, *SE* = 3.8) (**Figure 2C**). However, the change of RTs across the learning-set between trials in which the set-winner was chosen versus when the set-winner was not chosen was not statistically significant (interaction effect ‘RT by choosing set-winner versus not choosing set-winner’: *F*(2, 7829.6) = 0.082, *p* = 0.92).

**Figure 1.**
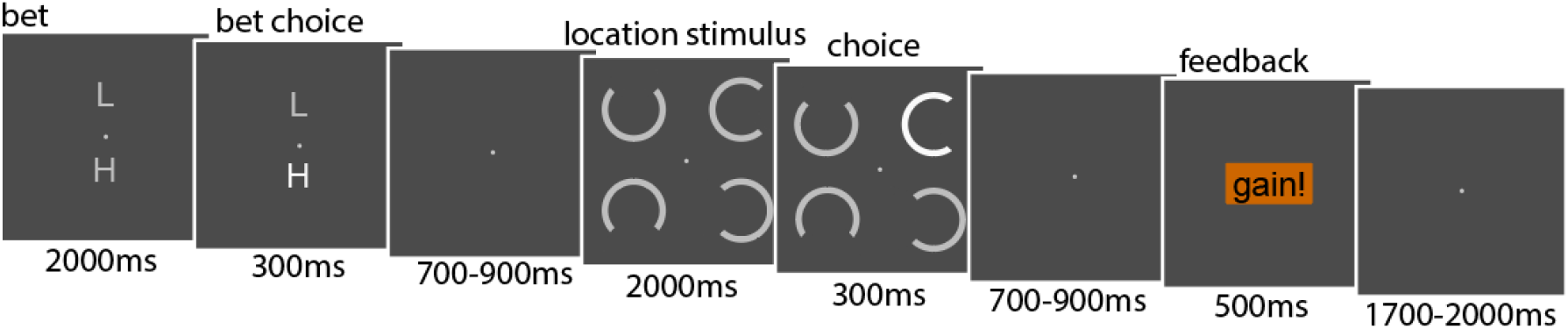
Trial-sequence of the probabilistic reward-based spatial learning task. First, participants had to choose between betting high or low by choosing the corresponding letter. Then, their choice of bet was highlighted. Next, participants chose a location by selecting the opening of the circle that was placed in the location on that trial. The choice of the opening of the circle was also highlighted after. Lastly, feedback in form of the words “gain!” and “loss!” appeared to guide their choice. Participants underwent the trial-sequence 40 times in each set before the set-winner changed.

**Figure 2.**
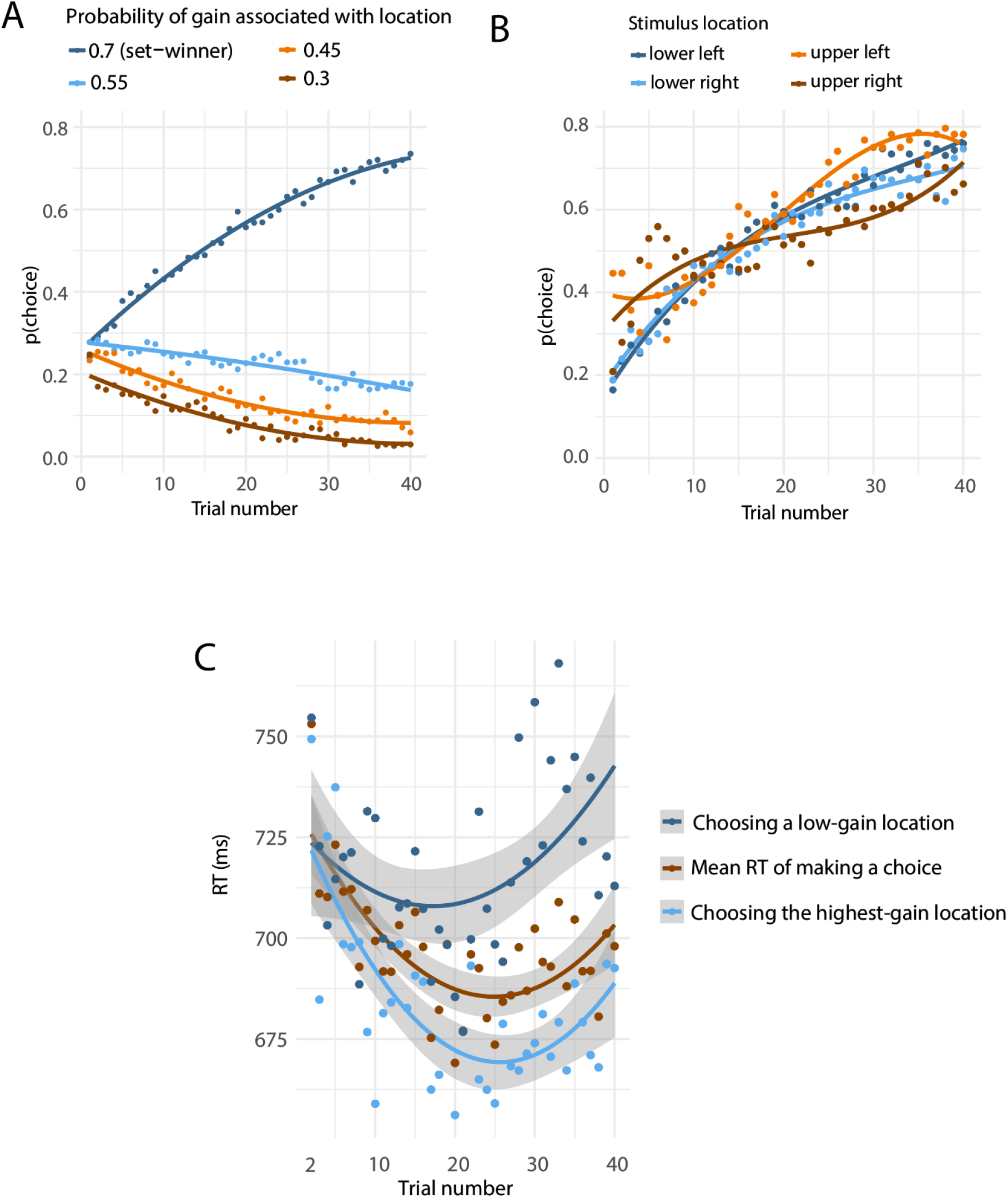
Behavioral results. (A) The probability of choosing a location with the highest gain probability increased with trial-number, whereas the probability of choosing all other locations decreased. (B) All four locations showed similar probability curves when they had the highest gain-probability assigned. (C) Participants were faster in choosing the highest-gain location-stimulus compared to the other location-stimuli.

### EEG

#### Feedback Processing

We observed a frontocentral ERP in the FRN latency range which amplitude was, as expected, significantly more negative in response to loss feedback compared to gain feedback (main effect ‘feedback’ *F*(1, 30) = 34.1, *p* < 0.001; *M*_gain-minus-loss_= - 1.38, *SE* = 0.24). However, we did not observe a statistically significant effect of trial-number on mean ERP amplitudes in the FRN latency range (main effect ‘trial-number’ *F*(1, 10190) = 1.25, *p* = 0.26), nor did we observe a statistically significant change of mean amplitudes with trial-number depending on loss or gain feedback (interaction effect ‘feedback by trial-number’, *F*(1, 10174) = 3.04, *p* = 0.08). Lastly, model comparison between a mixed model including an interaction effect of ‘feedback’ by ‘trial-number’ and a model without an interaction effect remained inconclusive based on the difference in AIC alone (AIC_Interaction_= 66121; AIC_Main effects only_= 66122). We therefore also took the BIC of both models into account during model selection, which revealed that the best fitted model was the more parsimonious main effects only model (BIC_main effects only_= 66172, BIC_Interaction_= 66179).

In line with our expectations, mean amplitudes in the LPC latency range showed significant changes with trial-number (main effect ‘trial-number’ *F*(1, 10180.3) = 9.34, *p* = 0.002). In addition, mean amplitudes in the LPC latency range significantly differed between gains and losses dependent on trial-number (interaction effect ‘trial-number by feedback’, *F*(1, 10172.9) = 19.82, *p* < 0.0001). As illustrated in **Figure 2B**, average amplitudes became more positive in response to losses with increasing trial-number, whereas gain-feedback elicited progressively smaller amplitudes (*M*_slope(gains)_= - 0.02, *SE* = 0.006; *M*_slope(losses)_= 0.02, *SE* = 0.007).

#### Alpha Power

As expected, we observed a significant post-feedback alpha suppression on occipital electrode sites contralateral to the upcoming location-stimulus of choice (*t*(28.35) = - 2.08, *p* = 0.046), supporting increased cortical activity in the visual cortex relative to the upcoming choice location (**Figure 4B and C**).

**Figure 3.**
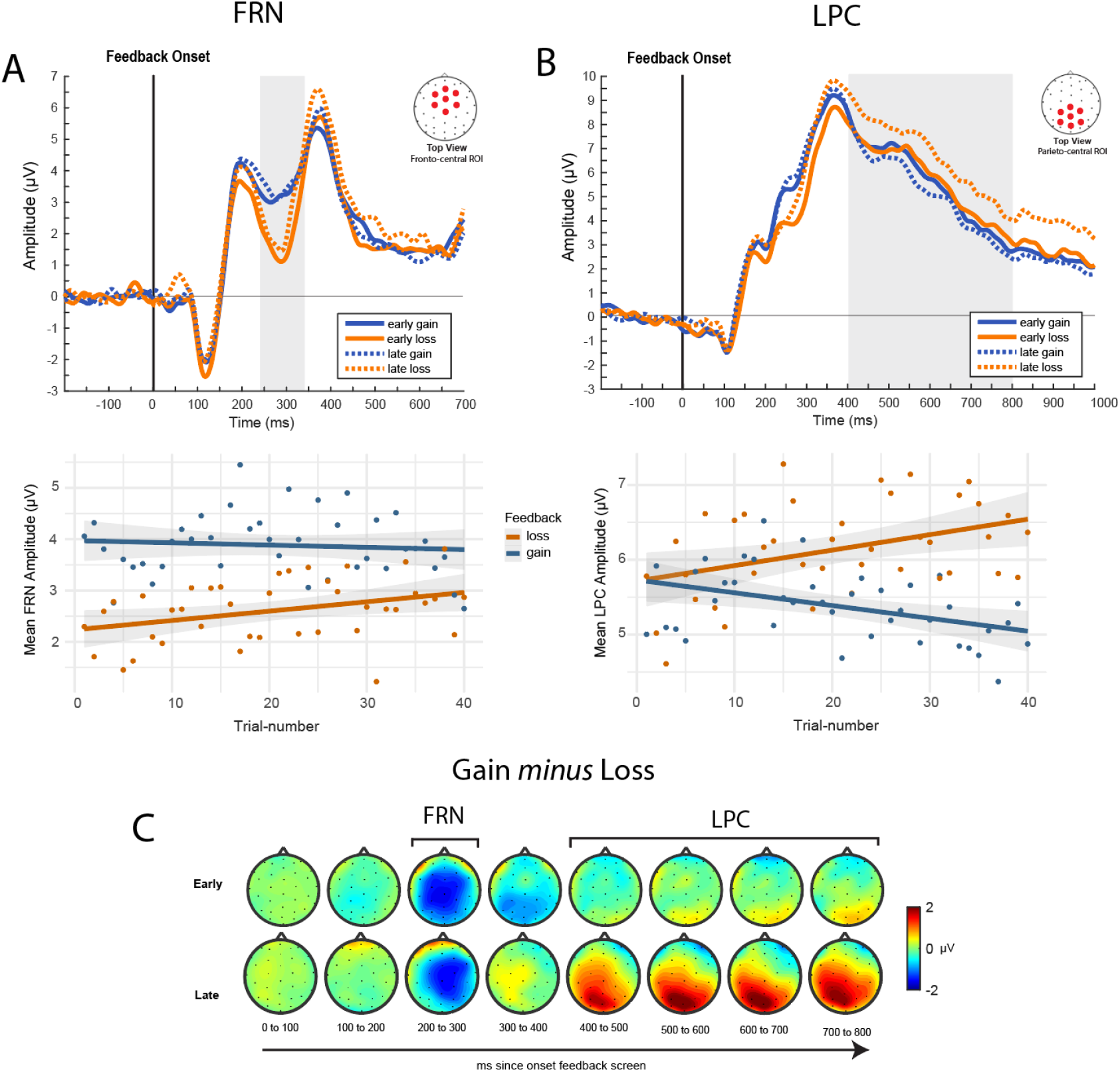
Feedback-locked ERPs early (first 8 trials) and late (last 8 trials) in the learning-set. (A) Fronto-central amplitudes between 240 and 340 ms were more negative in response to loss feedback, and showed no difference between early and late trials in the learning-set. (B) Parietal amplitudes between 400-800ms did not distinguish between losses and gains at the beginning of the learning-set, but shows larger responses to losses later on. (C) Topography distribution of the gain minus loss effect indicate a fronto-central FRN (200ms to 300ms) and a parieto-central LPC (400 to 800ms) early and late in the set. Values show the *difference* between gain and loss feedback (gain *minus* loss).

**Figure 4.**
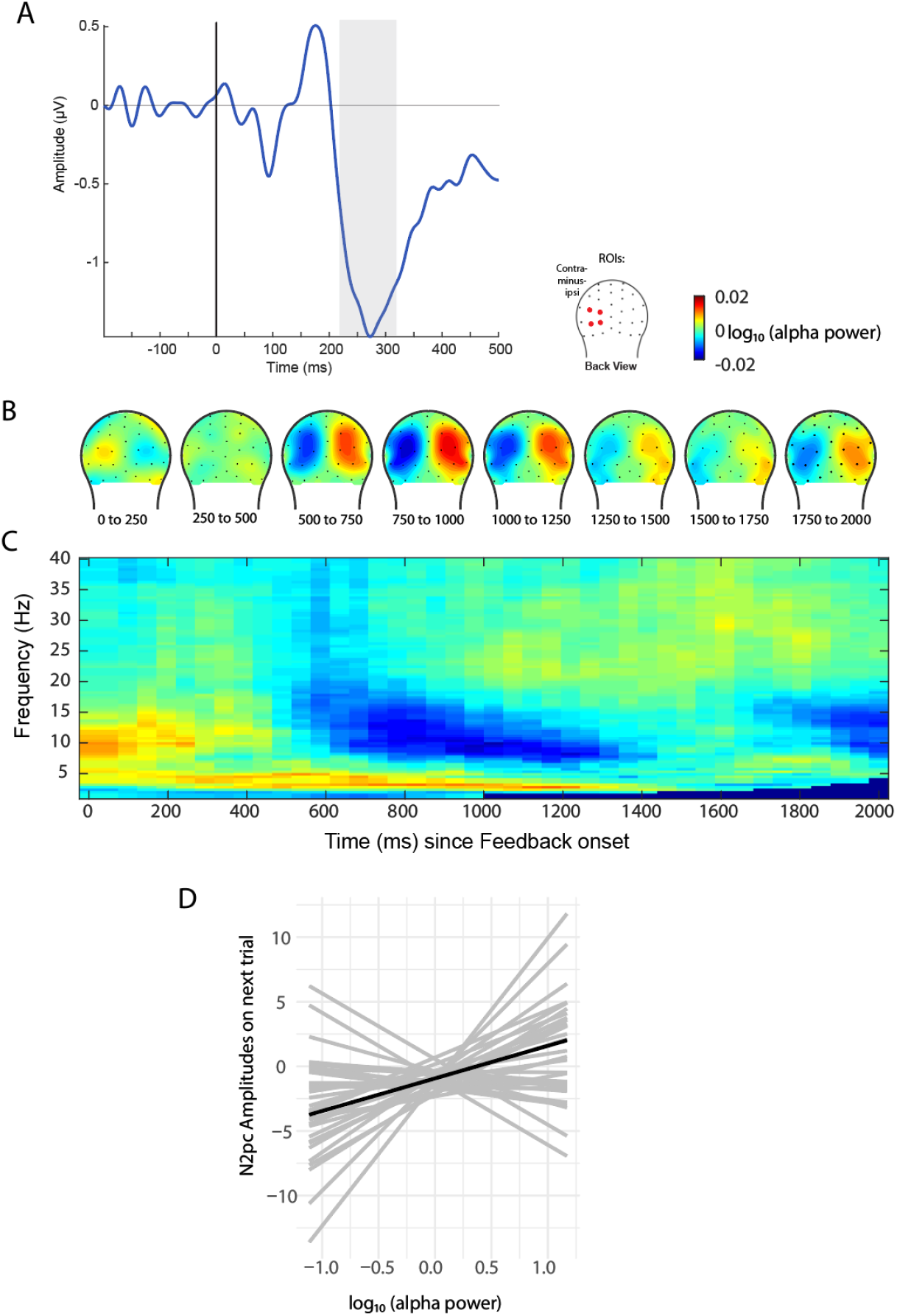
Neural attentional orienting in form of the N2pc component and preparatory bias in form of feedback-locked lateralized alpha power, calculated as a function of contralateral-versus-ipsilateral electrode sites to-choice. (A) Grand average N2pc trace relative to the location-stimulus of choice. We did not observe changes in mean choice-related N2pc amplitudes across the trial-set. (B) Topographical distribution of feedback-locked lateralized alpha power show that contralateral-to-upcoming choice alpha activity is significantly lower compared to ipsilateral activity at occipital electrode sites. (C) Time-frequency spectrum of our time-frequency analyses show that the decrease in contralateral power is specific for the alpha band (8-14Hz). (D) Individual subject’s regression slopes (grey) and mean regression slope (black) of the alpha-power versus N2pc analyses show that post-feedback lateralized-to-choice alpha power was significantly predictive of N2pc amplitudes on the next trial, in that the smaller the lateralized alpha power, the more negative N2pc amplitudes.

##### Relationship between alpha power and the N2pc

The presentation of the location-stimuli elicited an N2pc that was significantly larger on electrode sites that were contralateral to the stimulus-location of choice, reflecting the orienting of spatial attention towards the location of choice (*t*(30.18) = -7.06, *p* < 0.0001, **Figure 4A**). Next, we examined whether alpha power and N2pc amplitudes relative to the upcoming location choice would show a significant relationship. This was indeed the case (*F*(1, 28.02) = 5.04, *p* = 0.03). Specifically, the more pronounced suppression of post-feedback alpha power relative to the upcoming choice, the larger the N2pc amplitudes relative to the choice on the upcoming trial (**Figure 4D**).

## Discussion

Rewarded stimulus-features in our environment can voluntarily or involuntarily capture our attention (Anderson, 2016; Bourgeois et al., 2017). Yet, whether the same holds true for rewarded spatial locations is less straightforward (Sisk et al., 2020; Anderson & Kim, 2018, Chelazzi et al., 2014; Won & Leber, 2018). Here, we aimed to map out the cascade of neurocognitive mechanisms that underlie the learning of spatial stimulus-reward-associations in order to shed more light on the role of spatial reward-associations in value-based attentional capture. Our focus lied on neural reward-feedback processing, which is an essential part of learning (Schulz, 2015). We found significant differences in amplitudes between gain and loss feedback in the FRN and LPC, reflecting valence feedback and context-updating, respectively. Only the LPC showed significant modulations in amplitudes as the trial-set progressed. Subsequently, we observed a reduction in alpha power reflecting activity in the visual cortex contralateral to the upcoming location of choice, which was significantly related to the N2pc on the upcoming trial, a neural correlate for the orienting of spatial attention. Together, our findings provide insights into the neural cascades of neural reward feedback processing in the context of spatial attention.

By using a design with multiple unique learning sets of 40 trials, we were able to map the behavioral learning curves of spatial stimulus-reward associations. Similar behavioral results are also commonly reported in two-choice reward learning tasks which deploy non-spatial target-features as their to-be-rewarded stimuli (van den Berg et al., 2019; Bourgeois et al., 2017). In addition, the learning of stimulus-reward associations is a prerequisite in most studies investigating their attentional capture (Anderson et al., 2011; Anderson & Kim, 2018; Chelazzi et al., 2014). Our findings therefore replicate the behaviour that is found in feature-based attentional capture paradigms. Lastly, given that the average net gain was positive, these data support that participants actively used the feedback information to learn the spatial stimulus-reward association.

One of the prominent concepts in feedback processing is the reward prediction error (RPE), which reflects the difference in expected versus delivered feedback outcomes (Schultz, 2016). The FRN component has been associated with the RPE (Bellebaum & Daum, 2008, Nieuwenhuis et al., 2004, Cohen et al., 2007; Nieuwenhuis et al 2004). Given this interpretation, we expected to observe FRN amplitudes to be more negative for losses than for gains, and that this difference would become larger with trial-set progression, when the reward association is learned and reward expectancies become stronger. While we did observe FRN amplitudes to be different between loss and gain feedback, this difference did not change with trial-number, that is with learning. Our findings therefore suggest that the FRN represents feedback processing on a valence-level only, without playing a role in the RPE. Such a finding has been reported before (van Borries et al., 2013), and is further supported by our model comparison procedures, which showed that adding an interaction effect of feedback and trial-number did not improve model fit. Yet, it should be mentioned that, given that the highest-gain choice only delivered gains on a probability scale and participants were aware of that, loss feedback later in the set might not have been as unexpected. Work in recent years that challenges the FRN-RPE account theorized that it reflects a salience prediction error, reflecting the processing of informative versus uninformative outcomes (Talmi et al., 2013). Support for this view comes from findings that report larger FRN amplitudes for zero-value outcomes compared to loss or gain outcomes (Pfabigan et al., 2015; Huang & Yu, 2014). Taken together, while we did observe differences in FRN amplitudes between gain and loss feedback, we did not observe an effect of trial-number that would have supported the RPE account of the FRN.

Next to the FRN component, we also analysed amplitudes of the LPC in response to feedback and trial-number. The LPC has been suggested to reflect affective feedback processing and subsequent context-updating (Polich, 2007; Glazer et al., 2018). Against our expectations, we observed that, as the trial-set progressed, LPC amplitudes in response to loss feedback increased in size, whereas gain-feedback elicited gradually smaller LPC amplitudes. Late losses could signal that the current choice might need re-evaluation in order to stay on track of the most rewarding stimulus choice. This interpretation ties in with findings that report increased LPC amplitudes in response to unexpected losses (Trimber & Luhmann, 2017; Muller-Gras et al., 2019, Donaldson et al., 2016), and links between the LPC and subsequent behavioral adjustments (San Martín et al., 2013; Chase et al., 2011). Late gain feedback, on the other hand, does not provide any additional or new information about the context or the choice that is being made and no updating is needed, hence LPC amplitudes become smaller. Notably, research on the LPC in the context of learning from rewarding feedback is sparse, and the literature is just beginning to grow (Glazer et al., 2018).

Originally, the LPC is often associated with processes related to memory and affect (Hajcak et al., 2009; Kaller et al., 1995). Our findings do not exclude the interpretation that late loss feedback simply creates greater affective responses because the reward-association is known, but people still experience a loss, thereby eliciting greater responses in the LPC. Nonetheless, here we report distinct differences in LPC amplitudes in response to gain and loss feedback with learning, suggesting that LPC is dynamically modulated by learning stimulus-reward associations.

Following the modulation of the LPC, we observed that post-feedback alpha power was lower on occipital electrode sites contralateral to the upcoming choice. In general, power in the alpha band has been found to be inversely linked to the BOLD signal (Goldman et al., 2002; Scheeringa et al., 2011), wherein a reduction in alpha power is associated with increased cortical activity. Within this framework, our findings indicate increased activity in visual brain areas contralateral to the upcoming choice. Similar findings have been found in preparatory spatial attention, where brain activity read out through alpha power is interpreted as preparatory sensory activity for the to-be-attended stimulus location (Worden et al., 2000). Additionally, we observed that contralateral alpha power following feedback was predictive of N2pc amplitudes on the upcoming choice, suggesting that it could be related to subsequent attentional orienting to the next stimulus location. Specifically, preparatory activity in the sensory cortices is thought to facilitate subsequent attentional processing, as it is correlated with subsequent N2pc amplitudes relative to the attended location (Zhao et al., 2019), but also with decreased RTs (van den Berg et al., 2016). Multivariate pattern analyses also recently showed that preparatory contralateral alpha power boosts stimulus processing (Barne et al., 2020). Here, we extend those findings by reporting that preparatory alpha power also occurs in stimulus-reward learning tasks, as a direct function of feedback presentation.

Given the temporal order of neural feedback processing, disclosed by FRN and LPC, the outcome of feedback processing could lead to an “educated” decision on which choice should be made next. This, in turn, induces preparatory activity in the visual cortex related to the upcoming choice. This interpretation would be in line with results from invasive electrophysiological animal studies that have shown that neurons in the sensory cortices become tuned towards the reward-predicting stimulus as a result of reward learning, by showing more discriminability and excitability in response to it (Jurjut et al., 2017). Additional findings in mice have shown that visuospatial attention is accompanied by enhanced neuronal responses and synaptic activity in the visual cortex (Speed et al., 2020).

Few studies to date have investigated the role of the sensory cortices in reward feedback processing, specifically, the neurotemporal order of when they come into play. Neuroimaging studies demonstrated that the sensory cortices show activity post-feedback, but not *when* in the processing cascade this activation occurs (Schiffer et al., 2014), whereas invasive animal studies mainly focus on changes in neuronal firing patterns in the sensory cortices only, but not on cognitive processing that occurs beforehand and in higher-order brain areas (Jurjut et al., 2017; Speed et al., 2020). Lastly, one recent paper found that activity in the stimulus-specific sensory cortex occurred post-feedback of the *previous* choice (van den Berg et al., 2019). Our data suggests that activity in the visual cortex after feedback presentation is significantly related to subsequent attentional processing, and therefore could reflect preparatory mechanisms as a function of the feedback process.

While our findings show that spatial stimulus-reward associations can be learned and also affect subsequent attentional processing, one open question remains of whether the learned association can permanently capture attention in an unrelated task. Research on feature-based stimulus-reward-association learning suggests that reward-associations induce more permanent changes in the feature-related sensory cortex, which subsequently leads to attentional capture (Tankelevitch et al., 2020; MacLean & Giesbrecht, 2015). A similar proposition has been made for spatial stimulus-reward-associations (Chelazzi et al., 2014), but not yet been tested neurally. Our findings show temporary modulations in the visual cortex as a function of feedback processing, and future research could investigate whether and how reward-associations manifest themselves permanently in the sensory cortices.

To sum up, we set out to investigate the neuro-temporal cascade of feedback and spatial attentional processes that occur during spatial stimulus-reward learning. First, we observed that feedback valence processing as reflected through the FRN is not modulated by learning, and we conclude that the FRN only signals binary feedback outcome processing. The LPC, on the other hand, was dynamically modulated based on feedback and learning, supporting the view that it reflects neural context-updating. Next, we found that preparatory brain activity in the visual cortex at the beginning of each trial was lateralized towards the choice that was about to be made, and predictive of later attentional orienting read out via the N2pc. Our findings map out the temporal neural cascade of feedback processing that, ultimately, could lead to attentional capture by spatial reward associations.

